# Nutrient-Responsive Formation of Mitochondrial-Derived Structures in *Caenorhabditis elegans*

**DOI:** 10.1101/2025.06.24.661357

**Authors:** Miriam Valera-Alberni, Anne Lanjuin, Silvia Romero-Sanz, Kristopher Burkewitz, Adam L. Hughes, William B. Mair

**Affiliations:** Dept. Molecular Metabolism, Harvard TH Chan School of Public Health, Massachusetts, United States of America; Institute of Biomedicine and Molecular Genetics (IBGM), University of Valladolid–CSIC, Valladolid, Spain; Department of Biochemistry, Molecular Biology, and Physiology, Faculty of Medicine, University of Valladolid, Spain; Department of Cell and Developmental Biology, Vanderbilt University, Nashville, TN, United States of America; Department of Biochemistry, University of Utah School of Medicine, Salt Lake City, UT, United States of America

## Abstract

Mitochondrial morphology is dynamically regulated through remodeling processes essential for maintaining mitochondrial function and ensuring cellular and metabolic homeostasis. While classical models of mitochondrial dynamics center on cycles of fragmentation and elongation, emerging evidence highlights additional membrane remodeling mechanisms, including the formation of mitochondrial-derived vesicles (MDVs) and mitochondrial-derived compartments (MDCs). These mitochondrial-derived structures, however, have been predominantly characterized in cultured cells and unicellular organisms, leaving their relevance in multicellular systems largely unexplored. Here, we identify a previously uncharacterized class of mitochondrial-derived structures in *Caenorhabditis elegans* muscle cells that are induced in response to intermittent fasting. We show that these structures appear specifically during the refeeding phase— coinciding with mitochondrial elongation —and are absent during fasting. Consistent with MDCs, the structures, approximately 1 µm in size, are enriched in outer mitochondrial membrane markers such as TOMM-20^aa1-49^ and TOMM-70, but notably lack components of the inner mitochondrial membrane. Their formation requires the microtubule-associated MIRO-1/2 proteins, and their size is modulated by the mitochondrial dynamics machinery. Together, our findings reveal a nutritionally regulated mitochondrial remodeling event in *C. elegans* muscle that may play a role in mitochondrial quality control and adaptation to metabolic cues.

## Introduction

Mitochondria serve as central hubs of cellular metabolism, orchestrating a wide range of essential biological processes that sustain cell function and survival. While traditionally viewed as the primary producers of ATP, mitochondria also play crucial roles in fatty acid and heme biosynthesis, intracellular calcium homeostasis, thermogenesis, and the generation of reactive oxygen species (ROS) (Monzel et al., 2023). Metabolites and signaling intermediates produced within mitochondria, such as ROS, serve important regulatory functions and participate in signal transduction pathways (Chandel, 2014; Picard and Shirihai, 2022). Moreover, mitochondria maintain many contact sites with other organelles—including the endoplasmic reticulum, lipid droplets, endosomes, and peroxisomes— which support cellular homeostasis through dynamic communication events that regulate intracellular signaling, lipid metabolism, membrane dynamics, organelle division, and organelle biogenesis (Chen et al., 2025; Voeltz et al., 2024).

Mitochondrial shape is closely tied to function, as these organelles form dynamic networks that continuously remodel their morphology in response to physiological cues, adapting their architecture to meet metabolic demands and cellular stress (Liesa and Shirihai, 2013). This remodeling, termed mitochondrial dynamics, involves alternating states of elongation and fragmentation. Mitochondrial fission—the division of a mitochondrion into two daughter organelles—is mediated by a coordinated set of proteins, most notably the cytosolic GTPase dynamin-related protein 1 (DRP1), which is recruited to the outer mitochondrial membrane (OMM) to execute membrane scission. Conversely, mitochondrial fusion involves the merging of two mitochondria into one and is mediated in mammals by mitofusin 1 and 2 (MFN1 and MFN2) at the OMM and by optic atrophy 1 (OPA1) at the inner mitochondrial membrane (IMM). These core components of mitochondrial dynamics are evolutionarily conserved. In the case of the model organism *Caenorhabditis elegans*, mitochondrial fusion is regulated by FZO-1, the MFN1/2 homolog responsible for OMM fusion, and EAT-3, the OPA1 equivalent for inner membrane fusion, while mitochondrial fission is mediated by DRP-1 (Ichishita et al., 2008; Kanazawa et al., 2008; Labrousse et al., 1999; Rolland et al., 2009). Mitochondrial fission is recognized as an early step in mitophagy, facilitating the selective removal of damaged mitochondrial components by the autophagic machinery and thereby maintaining the integrity of the mitochondrial network (Twig et al., 2008). As such, mitochondrial dynamics serve as a quality control mechanism essential for cellular homeostasis. Disruption of this balance has been implicated in various pathological conditions, including metabolic disorders, neurodegenerative diseases, and cancer—conditions in which altered mitochondrial dynamics are closely linked to impaired mitochondrial function (Chan, 2020; Chen et al., 2023). Recent research has uncovered additional, non-canonical mechanisms of mitochondrial membrane remodeling. For instance, mitochondrial-derived vesicles (MDVs), originally identified for transporting cargo to peroxisomes, are small (60-150 nm) punctate structures that form in response to mild oxidative stress in mammalian cells and tissues (Braschi et al., 2010). Alternatively, elevated intracellular amino acid levels trigger the formation of large (1 µm) mitochondrial-derived compartments (MDCs) in yeast, which enable the selective removal of mitochondrial proteins, thereby facilitating cellular adaptation to amino acid excess (Schuler et al., 2021). Similarly, structures positive for the outer mitochondrial membrane (OMM), known as SPOTs, emerge during *Toxoplasma gondii* infection in mammalian cells (Li et al., 2022). Throughout this manuscript, we will collectively refer to these mechanisms of mitochondrial membrane remodeling as mitochondrial-derived structures. These structures reflect the growing complexity of mitochondrial membrane changes in response to nutrient imbalances and cellular stress signals. However, whether such structures also occur under physiological conditions in a multicellular animal remains unknown.

*C. elegans* provides a particular useful platform for visualizing mitochondrial morphology *in vivo*, due to its transparent cuticle, which permits noninvasive, high-resolution imaging of mitochondria across multiple tissues (Valera-Alberni *et al*, 2024). In this study, we report the discovery of a previously undescribed class of mitochondrial-derived structures that form in *C. elegans* in response to refeeding following periods of fasting. These structures are highly dynamic, appearing as mitochondrial spheres approximately 1 μm in diameter, and are enriched in outer membrane components, including the N-terminal domain of the translocase of the OMM, TOMM-20 (amino acids 1–49), or TOMM-70. Notably, they exclude inner mitochondrial membrane markers. The formation of these structures requires the conserved GTPase MIRO-1, but not the canonical mitochondrial dynamics proteins, which instead influence their size, findings that align with existing literature on mitochondrial-derived compartments (MDCs). Altogether, our findings describe a novel, nutrient-responsive mitochondrial remodeling event in *C. elegans*, expanding the current understanding of mitochondrial plasticity in response to metabolic cues.

## Results

### Formation of OMM-Enriched Mitochondrial Structures Upon Refeeding in *C. elegans*

Mitochondrial fusion and fission are integral processes for cellular and metabolic health. Many longevity promoting interventions require dynamic mitochondrial networks, such as intermitent fasting (IF) that lead to significant improvement of health and longevity across species (Longo et al., 2021). We have previously reported how mitochondrial networks in wild-type *C. elegans* animals undergo remodeling in response to IF, where networks cycle between fragmentation after fasting and elongation upon refeeding in muscle (Weir et al., 2017). To further delve into mitochondrial membrane remodeling associated with IF, we generated a *C. elegans* strain enabling dual labeling of mitochondrial membranes. The outer mitochondrial membrane (OMM) was labeled using a novel TOMM-20^aa1-49^::GFP construct driven by the *eft-3p* promoter, and the inner mitochondrial membrane (IMM) was labeled with an mScarlet tag fused to *sclp-4*, the *C. elegans* homolog of TIMM-50 (Valera-Alberni et al., 2024).

We next challenged the TOMM-20^aa1-49^::GFP; TIMM-50::mScarlet strain using an IF protocol in which animals are subjected to cycles of 2 days of fasting followed by 2 days of refeeding. In parallel, control animals were maintained on food *ad libitum* (AL) throughout the experiment. In these control worms, mitochondrial length progressively decreased with age in body wall muscle, reflecting an age-associated increase in mitochondrial fragmentation— a phenomenon widely reported across species (Lima et al., 2022; Liu et al., 2020). Contrarily, muscle mitochondria in the IF group were protected from excessive fragmentation due to the remarkable remodeling induced by the challenge. During fasting periods (days 4 and 8), mitochondrial fragmentation increased, as evidenced by a reduction in mitochondrial length **(Fig. 1, A and B)**. Upon refeeding (days 6 and 10), mitochondria underwent elongation, accompanied by a corresponding increase in mitochondrial length **(Fig. 1, A and B)**.

**Figure 1.**
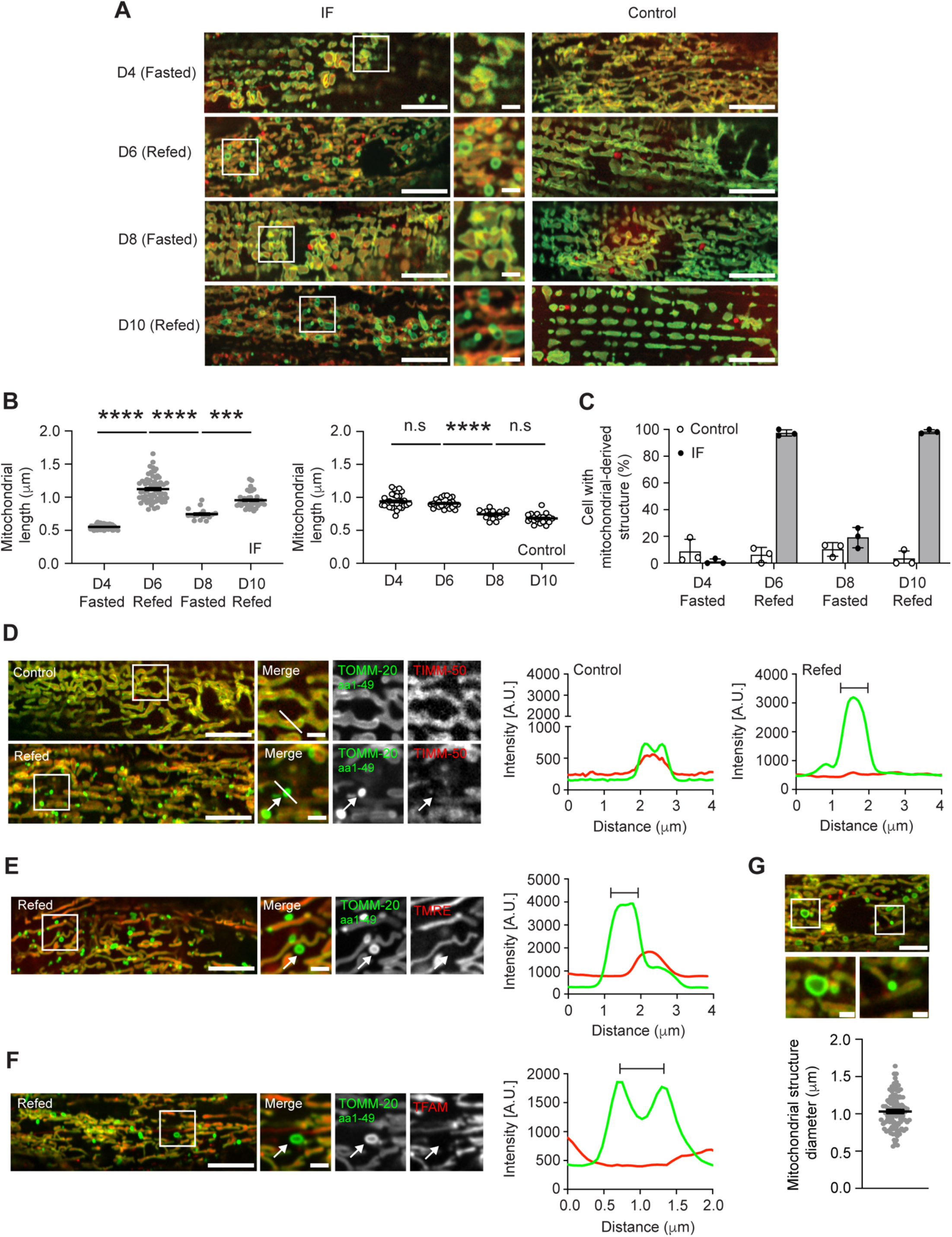
Mitochondrial-Derived Structures Form in *C. elegans* Muscle Following Refeeding. **(A)** Representative confocal fluorescence microscopy images of muscle cells from TOMM-20^aa1-49^::GFP; TIMM-50::mScarlet *C. elegans* demonstrate that mitochondrial networks undergo remodeling in response to intermittent fasting (IF), compared to control worms continuously kept fed. Scale bars, 10 µm and 2 µm**. (B)** Quantification of mitochondrial length shows a shift from fragmentation to elongation in IF worms (left), whereas control ad libitum-fed worms display an age-associated increase in mitochondrial fragmentation and a corresponding reduction in mitochondrial length (right). Data are presented as mean ± SEM. ***P < 0.001; ****P < 0.0001. **(C)** Quantification of the percentage of muscle cells containing at least one mitochondrial-derived structure for the indicated conditions using a TOMM-20^aa1-49^::GFP strain. Data are presented as mean ± SEM, with each data point representing a biological replicate. For each condition, 20–50 animals were analyzed. **(D)** Super-resolution confocal fluorescence microscopy images of muscle cells from TOMM-20^aa1-49^::GFP; TIMM-50::mScarlet worms at day 6 of adulthood, either maintained under continuous feeding (control) or refed after fasting. Scale bars, 10 µm and 2 µm. White line marks the position of the line-scan fluorescence intensity profile shown on the right. The bracket denotes the site of formation of the mitochondrial-derived structure. **(E)** Representative confocal fluorescence microscopy images of muscle from TOMM-20^aa1-49^::GFP refed worms stained with TMRE. Scale bars, 10 µm and 2 µm. **(F)** Representative confocal fluorescence microscopy images of muscle from TOMM-20^aa1-49^::GFP; HMG-5 (TFAM)::mScarlet refed worms. Scale bars, 10 µm and 2 µm. In D-F, the mitochondrial-derived structure is indicated by white arrows and the bracket in the intensity profile indicates the site of formation of the mitochondrial-derived structure. **(G)** Scatter plot showing the diameter of mitochondrial-derived structures in TOMM-20^aa1-49^::GFP; TIMM-50::mScarlet refed worms. Scale bars, 5 µm and 1 µm. Error bars represent the mean ± SEM of n = 100 MDCs from 10 different animals across 3 separate experiments.

Strikingly, upon refeeding at day 6 of adulthood, we observed the emergence of small, rounded structures connected to the main mitochondrial network that exhibited increased GFP intensity **(Fig. 1A)**. Notably, these mitochondrial-derived structures disappeared during subsequent fasting (day 8), when the network again became fragmented **(Fig. 1C)**. However, they reappeared upon refeeding at day 10, underscoring their dynamic and reversible nature in response to changes in the nutritional environment **(Fig. 1C)**.

Next, we sought to further characterize the properties of these *in vivo* induced mitochondrial-derived structures, and compare them to those previously defined for mitochondrial-derived vesicles (MDVs) or mitochondrial-derived compartments (MDCs). Line scan analysis revealed that the mitochondrial-derived structures induced upon refeeding are enriched nearly threefold in GFP intensity compared to the mitochondrial tubules to which they are associated **(Fig. 1D)**. Notably, these structures lack detectable TIMM-50::mScarlet signal, suggesting exclusion of the IMM **(Fig. 1D)**. Furthermore, they did not incorporate the mitochondrial membrane potential– dependent dye tetramethylrhodamine methyl ester (TMRE), indicating an absence of membrane potential and distinguishing them from the adjacent mitochondrial tubules, which were TMRE-positive **(Fig. 1E)**. To assess the presence of mitochondrial DNA (mtDNA), we generated a CRISPR knock-in strain expressing mScarlet fused to HMG-5, the *C. elegans* homolog of the mitochondrial transcription factor A (TFAM), a protein essential for mtDNA transcription, replication, and nucleoid organization (Bonekamp and Larsson, 2018; Brüser et al., 2021). When this strain was subjected to the fasting-refeeding challenge, the mitochondrial-derived structures clearly excluded TFAM, in contrast to the main mitochondrial network which included mScarlet signal, suggesting the *C.elegans* mitochondrial structures are devoid of mtDNA **(Fig. 1F)**. Collectively, these findings indicate that the mitochondrial-derived structures observed upon refeeding in *C. elegans* muscle are highly enriched in TOMM-20^aa1-49^::GFP but lack IMM components.

In our refeeding paradigm, we identified two distinct types of mitochondrial-derived structures: small, highly GFP-enriched puncta and larger, hollow ring-like structures **(Fig. 1G)**. Both consistently excluded TIMM-50::mScarlet and exhibited an average diameter of approximately 1 μm. An average diameter of approximately 1 µm—a relatively large size—is a characteristic feature of mitochondrial-derived compartments (MDCs), which are also enriched in the N-terminal fragment of TOMM-20 and lack IMM components. While mitochondrial-derived vesicles (MDVs) can also exclude the IMM, they are significantly smaller than MDCs. These observations provide an initial indication that the observed particles in *C. elegans* may represent MDCs.

### Molecular Cargo of Mitochondrial-Derived Structures in *C. elegans*

The different classes of mitochondrial-derived structures, including mitochondrial-derived vesicles (MDVs) and mitochondrial-derived compartments (MDCs), can be distinguished by their cargo composition. For instance, TOMM-20 is enriched in MDCs in yeast only when truncated to its N-terminal transmembrane domain (i.e., TOMM-20^aa1-49^), whereas the full-length TOMM-20 is not enriched (Wilson et al., 2024a). To characterize the molecular cargo of the mitochondrial-derived structures we observed in *C. elegans*, we first assessed whether full-length TOMM-20 localizes to these structures. To this end, we generated a strain expressing endogenously tagged TOMM-20::GFP. However, homozygous animals were sterile, and although heterozygotes were viable, their use was limited, as the tag clearly impacted endogenous TOMM-20 function and likely fails to accurately reflect whether it is incorporated into these structures.

To determine whether additional subunits of the TOMM complex are enriched in the *C. elegans* mitochondrial-derived structures, we next examined TOMM-70. Animals expressing endogenously tagged TOMM-70::GFP were viable, fertile, and phenotypically indistinguishable from wild-type controls, as we reported previously (Valera-Alberni et al., 2024). We thus analyzed the formation of mitochondrial-derived structures in these animals. Upon refeeding, we observed prominent formation of mitochondrial-derived structures enriched in TOMM-70::GFP in body wall muscle, which lacked TIMM-50::mScarlet **(Fig. 2A)**. These observations suggest that these novel *C.elegans* mitochondrial-derived structures could be enriched in multiple OMM components including TOMM-70.

**Figure 2.**
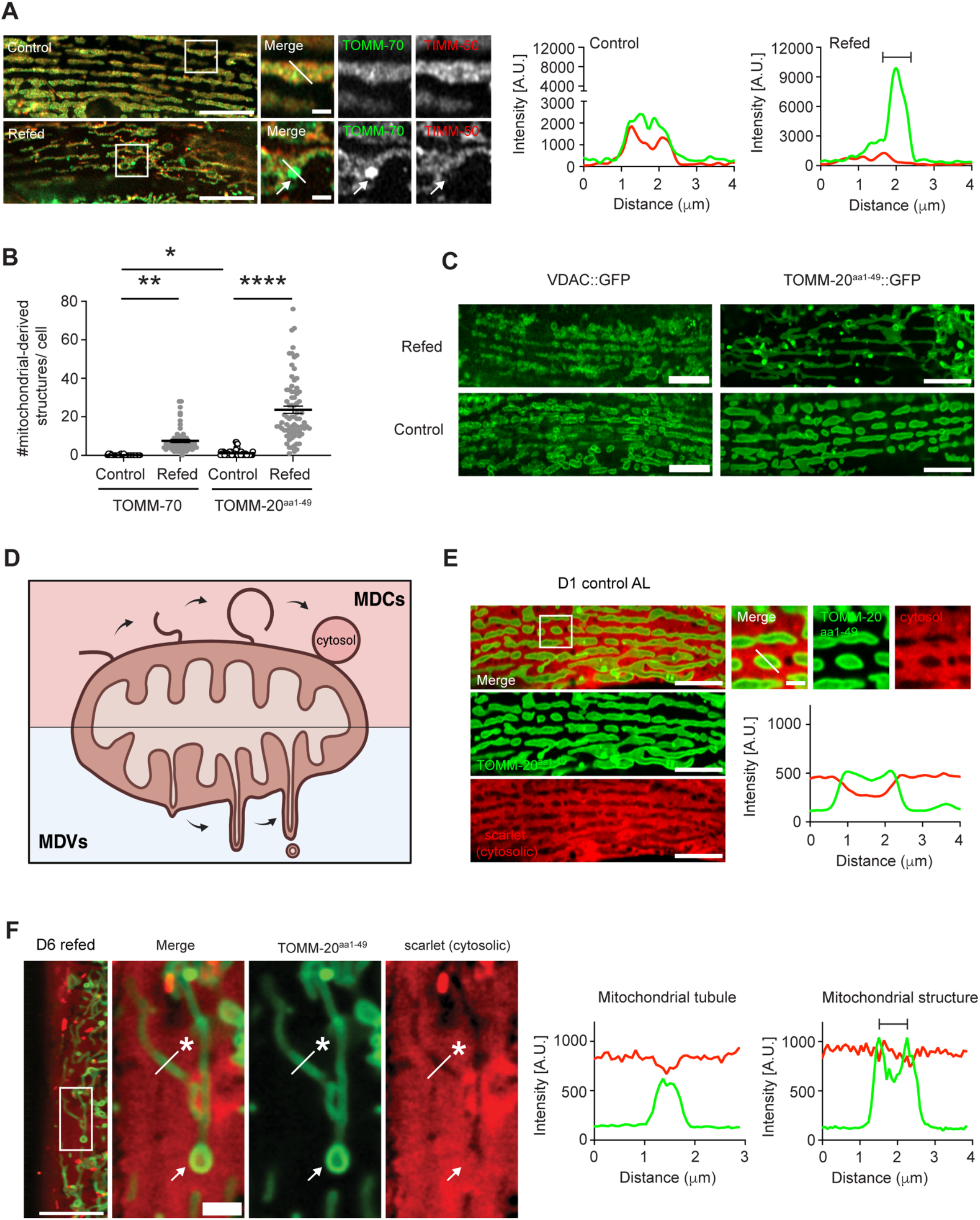
Molecular Identity and Characterization of Mitochondrial-Derived Structure Components. **(A)** Super-resolution confocal fluorescence microscopy images of muscle cells from TOMM-70::GFP; TIMM-50::mScarlet animals at day 6 of adulthood, either maintained under *ad libitum* (AL) feeding (control) or refed following an intermittent fasting protocol. Images were taken using an Airyscan detector. Scale bars, 10 µm and 2 µm. **(B)** Quantification of mitochondrial-derived structures per muscle cell in TOMM-70::GFP; TIMM-50::mScarlet and TOMM-20^aa1-49^::GFP; TIMM-50::mScarlet worms, respectively, at D6 of adulthood for control AL and refed animals. At least three experimental replicates were performed with n=30-50 worms per condition. All values are presented as mean ± SEM. *P < 0.05. **P < 0.01.****P < 0.0001. **(C)** Confocal fluorescence microscopy images of muscle cells from VDAC::GFP and TOMM-20^aa1-49^::GFP worms at day 6 of adulthood, comparing refed worms to AL animals. All images were obtained from the same experiment. Scale bars, 10 µm. **(D)** Schematic model illustrating the proposed biogenesis pathways of mitochondrial-derived compartments (MDCs) and mitochondrial-derived vesicles (MDVs). **(E-F)** Confocal fluorescence microscopy images of muscle cells from *myo-3p::tomm-20aa1-49::GFP::SL2::scarlet* animals at day 1 AL **(E)** or day 6 following refeeding **(F)**. The cytosolic mScarlet signal is excluded from the main mitochondrial tubules (white asterisk), except during structure formation, where mScarlet is incorporated (indicated with white arrow). Scale bars, 10 µm in E. Scale bars, 10 µm and 2 µm in F. Intensity profiles show GFP (green) and mScarlet (red) fluorescence signals. The bracket in the intensity profile indicates the site of formation of the mitochondrial-derived structure.

Quantification of the newly formed mitochondrial-derived structures revealed a significant increase in their number upon refeeding in both TOMM-70::GFP and TOMM-20^aa1-49^::GFP animals, compared to their respective AL-fed controls **(Fig. 2B)**. Notably, TOMM-20^aa1-49^::GFP animals exhibited a markedly higher number of structures following refeeding (23.66 ± 2.114 structures per muscle cell) compared to refed TOMM-70::GFP animals (7.517 ± 1.058 structures per muscle cell), suggesting that overexpression of TOMM-20^aa1-49^ may enhance the formation of mitochondrial-derived structures. Supporting this observation, even comparing AL conditions, TOMM-20^aa1-49^::GFP animals displayed significantly more mitochondrial-derived structures than TOMM-70::GFP animals **(Fig. 2B)**. Importantly, a wide range of OMM proteins have been reported to induce the formation of mitochondrial-derived compartments (MDCs) in yeast, when individually overexpressed (Wilson et al., 2024a). However, not all OMM proteins are incorporated into MDCs to the same extent. Core subunits of the TOM complex are generally excluded from MDCs unless the assembly of the complex is disrupted, thus explaining why MDCs include the TOMM-20^aa1-49^ domain but not the full-length protein (Wilson et al., 2024a). Notably, MDCs are also de-enriched of the voltage-dependent anion channel (VDAC, known as Por1 in yeast) (Wilson et al., 2024a). Consistent with this, *C.elegans* animals expressing VDAC tagged with GFP failed to form detectable mitochondrial-derived structures upon refeeding, in contrast to TOMM-20^aa1-49^::GFP animals analyzed in parallel, which did **(Fig. 2C**). In fact, the exclusion of β-barrel proteins—including TOMM-40, SAMM-50, and VDAC—is a defining feature that distinguishes MDCs from mitochondrial-derived vesicles (MDVs) (König et al., 2021). While VDAC is highly enriched in TOMM-positive MDVs, it is excluded from MDCs.

MDVs and MDCs also differ in their biogenesis mechanisms **(Fig. 2D)**. MDV formation involves membrane curvature, tubulation, and scission, whereas MDCs are formed through an OMM extension that rounds and encloses cytosolic content (König and McBride, 2024; Wilson et al., 2024b). To test whether the mitochondrial-derived structures in *C. elegans* incorporate cytosolic material, we generated a strain in which TOMM-20^aa1-49^::GFP and cytosolic mScarlet are co-expressed in muscle cells via a bicistronic cassette **(Fig. 2E)**. In these cells, mScarlet filled the entire cytosol and nucleoplasm but was clearly excluded from the interior of mitochondrial tubules **(Fig. 2E)**. After subjecting these worms to the fasting-refeeding challenge, the mScarlet signal remained excluded from the mitochondrial tubules (asterisk, **Fig. 2F**). However, the cytosolic mScarlet signal was detected inside the mitochondrial-derived structures that appeared upon refeeding (arrow, **Fig. 2E**), supporting the notion that these structures may entrap cytosolic material in a manner consistent with MDCs.

### Temporal Dynamics of Mitochondrial Structure Formation

To determine the temporal dynamics of structure formation during refeeding, we performed a time-course imaging experiment, capturing images hourly from the onset of refeeding after animals had been fasted for 48 hours. Notably, mitochondrial-derived structures did not form immediately after placing the worms on food but instead appeared in conjunction with the elongation of the mitochondrial network. During the first hour of refeeding, mitochondria remained fragmented **(Fig. 3A)**. However, by 2 hours, we observed a transition to a more elongated mitochondrial network, accompanied by the initial appearance of mitochondrial-derived structures **(Fig. 3A)**. These included both partially formed structures (**Fig. 3B**, upper inset) and fully formed, circular structures (**Fig. 3B**, lower inset). These observations suggest that the formation of mitochondrial-derived structures is closely linked to the elongation of the specific mitochondrial tubule they are associated with. The number of these structures progressively increased over time, reaching an average of 17 per muscle cell by 8 hours **(Fig. 3C)**. Interestingly, our previous data show that these structures remain detectable even after 48 hours of refeeding **(Fig. 1D)**, raising the question of whether they are stably maintained or continuously recycled during this period.

**Figure 3.**
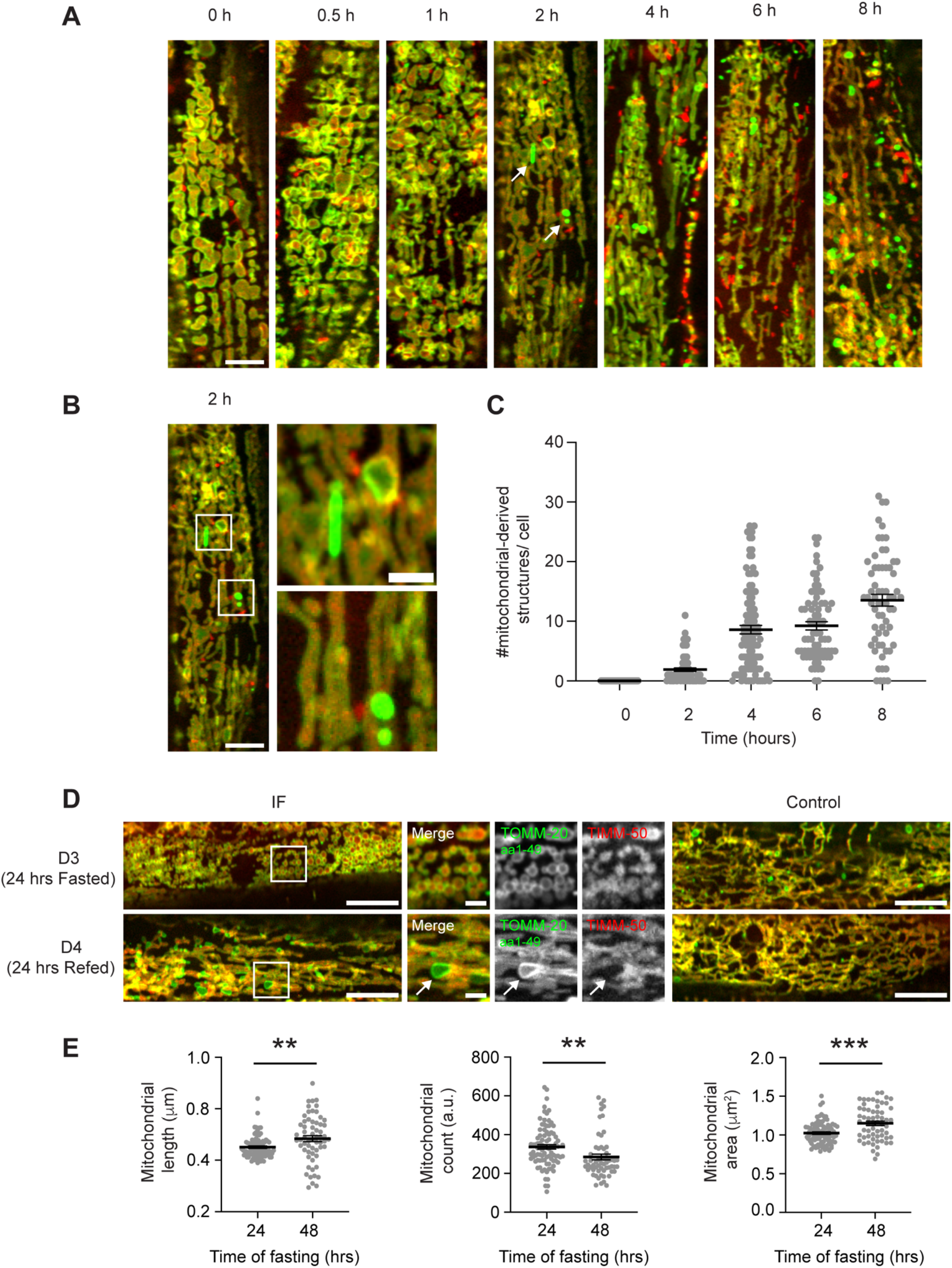
Temporal Dynamics of Mitochondrial-Derived Structure Formation. **(A)** Representative confocal fluorescence images of muscle cells from *C. elegans* TOMM-20^aa1-49^::GFP; TIMM-50::mScarlet worms during a refeeding time course. The 0 h time point corresponds to the end of a 48-hour fasting period, while subsequent time points show mitochondria after refeeding. White arrows at 2 h indicate the appearance of mitochondrial-derived structures. Scale bar, 10 µm. **(B)** Higher magnification views showing details of mitochondrial-derived structure formation at 2 h post-refeeding. Scale bars, 10 µm and 2 µm. **(C)** Quantification of the number of mitochondrial-derived structures per muscle cell at the indicated time points. Data represent mean ± SEM from n = 20–30 animals across 3 independent experiments. **(D)** Representative confocal images of muscle cells from TOMM-20^aa1-49^::GFP; TIMM-50::mScarlet animals following a 24-hour intermittent fasting paradigm. Animals were fasted for 24 h (Day 3, fasted) and then refed for 24 h (Day 4, refed). Images from age-matched animals maintained on *ad libitum* (AL) food are shown for comparison. Scale bars, 10 µm (overview) and 2 µm (insets). **(E)** Quantification of mitochondrial length, number, and area of mitochondria per muscle cell in animals subjected to 24 h or 48 h fasting. Data represent mean ± SEM from n = 20–30 animals across 3 independent experiments. **P < 0.01, ***P < 0.001.

In our protocol, we observed the formation of mitochondrial-derived structures in *C. elegans* muscle following a 48-hour intermittent fasting regimen, which has been reported to promote healthspan and lifespan in *C. elegans* (Honjoh et al., 2009; Uno et al., 2013; Weir et al., 2017). However, since 48 hours of fasting is a prolonged period relative to the lifespan of *C. elegans*, we investigated whether a shorter fasting regimen could similarly induce the formation of these structures. To this end, we implemented an alternate-day fasting protocol, consisting of 24 hours of fasting followed by 24 hours of refeeding. Mitochondria in fasted animals appeared fragmented (**Fig. 3D**), although fragmentation was less pronounced than that observed after 48 hours of fasting (**Fig. 3A** and **3D**). In fact, mitochondria following 48 hours of fasting were fragmented yet larger in size compared to those after 24 hours of fasting, as indicated by a reduced mitochondrial count and increased average mitochondrial area (**Fig. 3E**).

Although mitochondrial-derived structures were detectable following the 24-hour fasting routine, many appeared incompletely formed (**Fig. 3D**). We frequently observed TOMM-20^aa1-49^::GFP-enriched regions budding from the mitochondrial tubules, rather than fully separated structures (**Fig. 3D**). Whether this reflects impaired formation or simply indicates that more time is required for full maturation remains unclear. Further studies will be needed to elucidate the mechanisms governing their formation under different fasting conditions.

### Regulation of Mitochondrial Structure Formation in *C. elegans*

Over the past years, substantial efforts have been dedicated to elucidating the molecular machinery responsible for the formation of the different mitochondrial-derived structures, including MDVs, MDCs, and related membrane remodelling forms. Despite the varying nomenclature used to describe these structures, these processes generally follow common steps found in vesicle transport pathways including initiation, where cargo is selected and vesicle formation is initiated; tubulation, involving membrane protrusion or elongation; and scission, where the vesicle is separated from the mitochondrion (König and McBride, 2024; Wilson et al., 2024b). With this framework in mind, we next sought to investigate the molecular components driving the formation of the mitochondrial-derived structures observed in *C. elegans*.

Microtubule-dependent pulling forces have been shown to contribute to the formation of both MDVs and MDCs by extending thin membrane tubules from mitochondria, a process mediated by the OMM Rho GTPases MIRO1 and MIRO2. These calcium-regulated GTPases anchor mitochondria to motor proteins of the kinesin and dynein families (Canty et al., 2023; Eberhardt et al., 2020). To test their role in *C. elegans*, we used an RNAi construct that targets both *miro-1* and *miro-2*. As previously reported, *miro-1/2* RNAi induced the formation of thread-like mitochondrial tubules **(Fig 4A)**, consistent with phenotypes observed in *miro-1* deletion mutants (Ren et al., 2023). Notably, *miro-1/2* RNAi significantly reduced the formation of mitochondrial-derived structures under refeeding conditions compared to EV RNAi **(Fig 4A and 4B)**, implicating MIRO-1/2 in the remodeling events leading to the *C.elegans* mitochondrial-derived structure formation.

**Figure 4.**
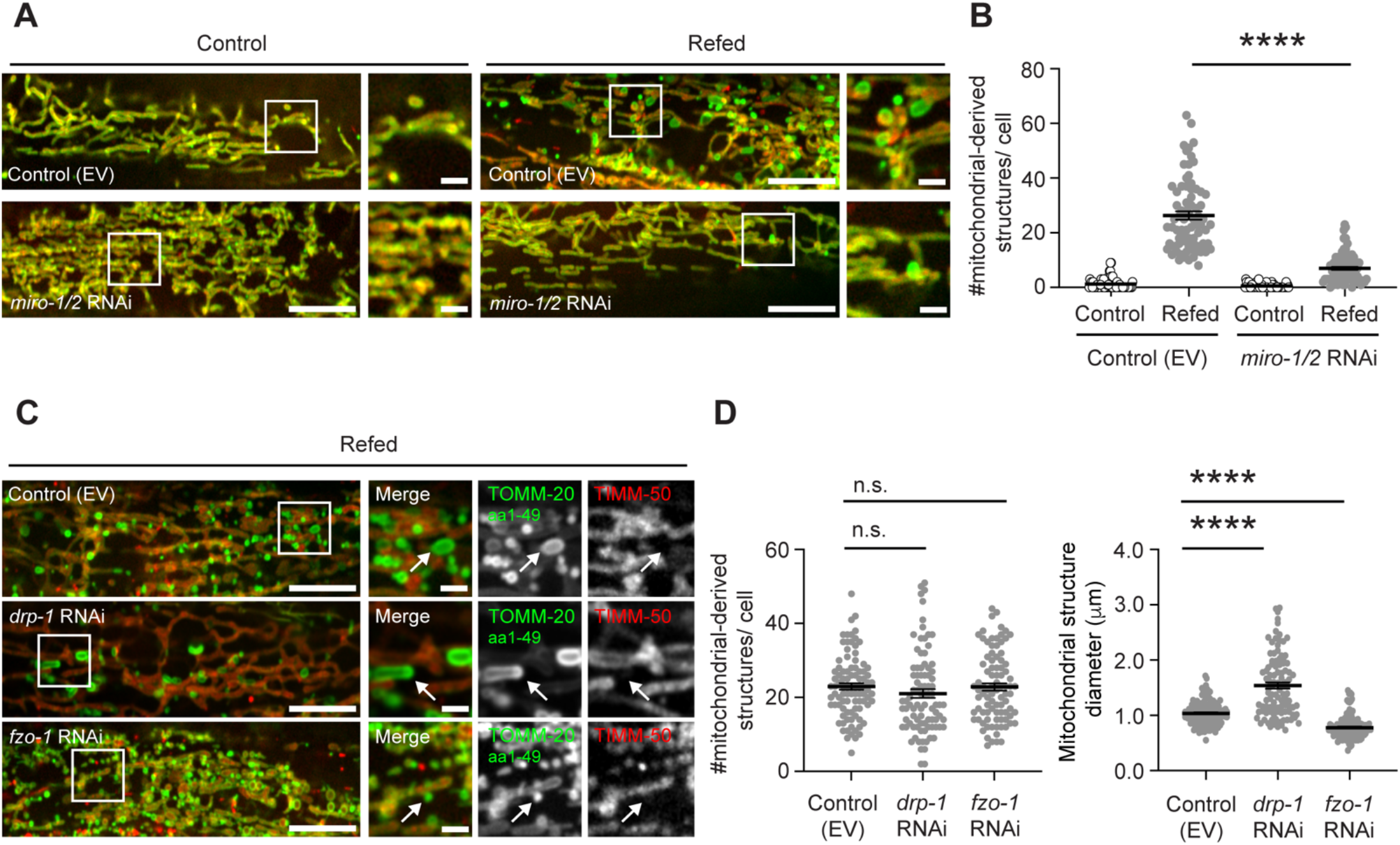
Modulators of Mitochondrial-Derived Structure Formation. **(A–D)** Representative confocal fluorescence images of muscle cells from *C. elegans* expressing TOMM-20^aa1-49^::GFP; TIMM-50::mScarlet at day 6 of adulthood. Animals were either maintained on *ad libitum* (AL) feeding (control) or subjected to 48 hours of fasting followed by 48 hours of refeeding. RNAi treatments targeting specific genes were used to assess their role in mitochondrial-derived structure formation and were compared to an empty vector (EV) RNAi control. **(A)** Analysis of the mitochondrial trafficking regulators *miro-1* and *miro-2*. **(B)** Quantification of mitochondrial-derived structures upon *miro-1/2* RNAi. **(C)** Disruption of mitochondrial dynamics via RNAi against *drp-1* (fission) and *fzo-1* (fusion). **(D)** Quantification of both mitochondrial-derived structure number and structure diameter in response to *drp-1* and *fzo-1* RNAi. Data represent mean ± SEM from three independent experiments. ****P < 0.0001. Scale bars: 10 µm (overview) and 2 µm (insets).

While MIRO-driven tubule elongation is a shared feature of both MDCs and MDVs, a key distinguishing factor is the requirement for DRP1, a key GTPase involved in mitochondrial fission. In MDVs, DRP1 localizes to the tips of membrane protrusions, where it facilitates vesicle scission; loss of DRP1 leads to an almost complete inhibition of TOMM-20–positive MDV formation in COS-7 cells (König et al., 2021). In contrast, DRP1 is not required for MDC formation (Hughes et al., 2016; Wilson et al., 2024b). Therefore, we next tested the role of DRP1 in the formation of the mitochondrial-derived structures in *C.elegans*. In control animals, RNAi-mediated knockdown of *drp-1* resulted in a pronounced elongation of the mitochondrial network compared to animals treated with no RNAi **(Suppl Fig 1A)**. Conversely, RNAi targeting *fzo-1*—the sole *C. elegan*s homolog of the mammalian MFN1/2—induced a prominent shift toward a fragmented mitochondrial morphology **(Suppl Fig 1A)**. Upon refeeding, mitochondrial-derived structures were still observed in *drp-1* RNAi-treated worms, indicating that their formation does not require DRP1 **(Fig 4C)**. Interestingly, these structures often appeared larger and were more distinctly arranged into extended domains compared to controls **(Fig 4C and 4D)**. In contrast, RNAi specific to *fzo-1* resulted in the accumulation of smaller TOMM-20^aa1-49^::GFP – enriched puncta that excluded TIMM-50::mScarlet **(Fig 4C and 4D)**. These changes in mitochondrial-derived structure size following *drp-1* and *fzo-1* RNAi were not accompanied by changes in mitochondrial-derived structure number, which remained comparable to the control (EV) **(Fig 4D)**. These results support the idea that mitochondrial elongation facilitates the early steps of structure formation, producing protrusions approximately 1 μm in size under control conditions. DRP1 may then act to constrict or sever outer membrane extensions. Importantly, the formation of TOMM-20^aa1-49^::GFP –enriched structures in *drp-1* RNAi animals suggests that the structures we observe are more reminiscent of MDCs, which do not require DRP1 for their formation.

In summary, these findings provide the first characterization of the formation of mitochondrial-derived structures in the multicellular animal *C. elegans*, revealing features—such as size, molecular cargo, and regulatory mechanisms of formation—that are consistent with mitochondrial-derived compartments (MDCs). This suggests that MDC-like remodeling events are conserved across species. Importantly, we show that such mitochondrial remodeling occurs *in vivo* in response to refeeding, a physiologically and ecologically relevant cue. While the precise function of these structures and their full set of regulators remain to be defined, our study represents a critical step toward understanding how mitochondrial architecture adapts to nutrient availability in a living animal.

## Discussion

Mitochondria are central hubs of cellular metabolism and play a critical role in coordinating intracellular signaling pathways. Their ability to undergo dynamic processes—such as fusion, fission, and interactions with other organelles—directly influences mitochondrial function. These structural adaptations are highly responsive to environmental cues, including nutrient availability and cellular stress, thus functioning as essential quality control mechanisms that regulate cell fate, metabolic homeostasis, and disease progression. Since the early 20th century, when scientists like M.R. Lewis first documented mitochondrial morphological changes (Lewis and Lewis, 1914), advances in high-resolution imaging have significantly deepened our understanding of mitochondrial membrane remodeling mechanisms. In this study, we report the discovery of nutrient-dependent, mitochondrial-derived structures in *Caenorhabditis elegans*, which emerge in response to refeeding following an intermittent fasting (IF) regimen.

Intermittent fasting, characterized by alternating periods of food deprivation and refeeding, has emerged as a promising strategy to promote healthy aging and metabolic resilience across species. In *C. elegans*, we previously showed that IF leads to marked changes in mitochondrial architecture, characterized by increased fragmentation following fasting and a rebound in network elongation upon refeeding (Weir et al., 2017). In the current study, we observed that this refeeding phase coincides with the appearance of distinct mitochondrial-derived structures in body wall muscle cells. These structures emerge approximately two hours after food reintroduction, temporally aligned with the initiation of mitochondrial elongation, and are notably absent during the fragmented state induced by renewed fasting. This correlation suggests that mitochondrial elongation may be a prerequisite for the formation of these structures, although the underlying molecular mechanisms remain to be elucidated. To investigate the factors driving the formation of these structures, it would be valuable to examine the changes in nutrient and metabolite levels that occur in *C. elegans* upon refeeding, and to determine whether the formation of mitochondrial-derived structures is linked to a specific nutrient cue. For instance, amino acid overload has been associated with the formation of a particular form of mitochondrial-derived structures, mitochondrial-derived comparmtents (MDCs), in yeast. MDCs are induced upon acute pharmacological inhibition of vacuolar acidification using concanamycin A (ConcA), a specific inhibitor of the evolutionarily conserved vacuolar H^+^ -ATPase (V-ATPase) proton pump (Schuler et al., 2021). In fact, we conducted preliminary experiments treating *C. elegans* with Concanamycin A following established protocols (Wei and Ruvkun, 2020), and observed a mild increase in the number of mitochondrial-derived structures **(Suppl Fig 2A)**. However, factors such as compound bioavailability, uptake efficiency, and treatment duration likely influence these outcomes, and further optimization is warranted. In yeast, MDCs are also known to form during aging (Hughes et al., 2016). Interestingly, in our study, we detected an increased number of mitochondrial-derived structures following the second cycle of refeeding at day 10, compared to the first cycle at day 6 **(Suppl Fig 2B)**, suggesting that aging may similarly promote the formation of these structures in *C. elegans*.

*C. elegans* offers unique advantages for investigating mitochondrial morphology *in vivo*, including its transparent cuticle, well-defined tissue organization, and amenability to live imaging without fixation—an important consideration, given that fixation procedures can artifactually alter mitochondrial-derived structures. This has been observed in the case of MDCs, where fixation and permeabilization in cultured cells causes TOMM70-enriched structures to collapse into smaller puncta lacking a visible lumen (Schuler et al., 2020). In contrast, *C. elegans* offers the advantage of non-invasive imaging, allowing for the preservation of cellular structures and dynamics within a physiological context. Mitochondria from different tissues exhibit distinct fuel preferences and biosynthetic roles (Vafai & Mootha, 2012), resulting in tissue-specific differences in mitochondrial content and morphology. This is also evident in *C. elegans*, where the intestine shows a higher degree of mitochondrial fragmentation, while the hypodermis exhibits prominent mitochondrial elongation (Valera-Alberni et al., 2024). In comparison, the body wall muscle displays a more balanced state between fragmentation and elongation, and was therefore selected as the primary tissue for this study. Specifically in muscle, we observe the formation of mitochondrial-derived structures associated with the shift in mitochondrial morphology in response to intermittent fasting, emerging upon refeeding when mitochondria transition to a more elongated state. Whether similar structures form in other tissues during physiological and nutrient adaptation remains to be determined.

With the discovery of new mitochondrial membrane remodeling pathways, a central question is whether the mitochondrial formations we observe in *C. elegans* align with known mitochondrial-derived structures, such as MDCs or mitochondrial-derived vesicles (MDVs). In our study, we identify large, circular mitochondrial-derived structures enriched in TOMM-70 but lacking VDAC and IMM components— features more consistent with MDCs than MDVs (König et al., 2021; König and McBride, 2024). A primary distinction between MDCs and MDVs lies in size: MDVs are small vesicles (60–150 nm), whereas MDCs are larger (typically around 1 µm). Our *C. elegans* structures average approximately 1 µm in diameter, supporting their alignment with MDCs. Another difference involves cargo: while a specific class of MDVs exclude matrix and IMM proteins (e.g., TOMM-20+/PDH−), these TOMM-20+-MDVs retain most cellular β-barrel proteins, such as VDAC. In contrast, MDCs do not incorporate full-length TOMM-20 or VDAC. Instead, MDCs selectively enrich the N-terminal domain TOMM-20^aa1-49^ (Wilson et al., 2024a). Consistent with this, our *C. elegans* structures show an enrichment in TOMM-20^aa1-49^, while being depleted in VDAC. Moreover, they exclude IMM proteins testified by the exclusion of TIMM-50, lack of mtDNA and mitochondrial membrane potential. We attempted to define whether the full-length TOMM-20 could be enriched in *C. elegans* structures using an endogenous full-length TOMM-20::GFP reporter, but developmental abnormalities in this strain limited interpretation. Further studies are needed to clarify whether the full-length TOMM-20 is present in the structures we observe in worms. Nonetheless, the molecular and structural features of these compartments strongly support the conclusion that the mitochondrial-derived structures we observe in *C. elegans* are bona fide mitochondrial-derived compartments (MDCs).

Moreover, the formation mechanisms of the mitochondrial-derived structures we observe in *C. elegans* closely resemble those reported for MDCs in yeast. MDCs originate through elongation and invagination of the OMM, sequestering cytosolic material (**Fig. 2D**) (Wilson et al., 2024b). Moreover, their formation depends on Gem1 in yeast (MIRO in mammalian cells), but occurs independently of DRP1 (English et al., 2020; Wilson et al., 2024b). Our data indicate that *C. elegans* structures may sequester cytosolic content and form independently of DRP1. Instead, downregulation of *drp-1* results in the appearance of larger, irregularly shaped structures, suggesting that DRP1 may act at a late stage to constrict OMM protrusions, facilitating their maturation into spherical compartments. Furthermore, we find that their formation relies on MIRO-1/2. Taken together, our findings suggest that the *C. elegans* structures resemble MDCs more closely than MDVs, although they may share overlapping molecular machinery. Future work should aim to more clearly define the boundaries between these pathways or determine whether they represent points along a shared mitochondrial quality control continuum, modulated by stress type or timing. This distinction is crucial for advancing our understanding of mitochondrial homeostasis.

In addition to elucidating the core mechanisms underlying the formation of these *C.elegans* mitochondrial-derived structures, understanding their functional relevance and identifying their target organelle(s) is equally important. A key outstanding question is: what is their purpose? Answering this will require uncovering their ultimate destination through detailed spatial and temporal analyses, including investigation of potential interactions with specific organelles. In other systems, MDVs are known to deliver cargo to peroxisomes for functional integration or *de novo* biogenesis, or to lysosomes and multivesicular bodies as part of mitochondrial quality control signaling (König and McBride, 2024). In parallel, MDCs form at ER–mitochondria contact sites and persist at these locations before being trafficked to lysosomes (or vacuoles in yeast) where the enclosed material is degraded (English et al., 2020). It will therefore be important to examine whether the *C. elegans* mitochondrial-derived structures associate with any of these organelles. Interestingly, when a second fasting cycle was applied to previously refed worms harboring these structures, we observed the expected shift toward fragmentation of the main mitochondrial network, accompanied by the disappearance of most mitochondrial-derived structures. Whether this disappearance reflects reintegration into the mitochondria, targeting to another organelle, or degradation via lysosomes remains to be determined. Finally, understanding how these events are integrated into the organism’s physiological response to intermittent fasting will be essential. Elucidating the functional role of these structures in the broader context of nutrient sensing and metabolic remodeling could uncover novel aspects of mitochondrial biology and open the door to many exciting discoveries.

In summary, our data indicate that the structures observed in *C. elegans* correspond to mitochondrial-derived compartments (MDCs). This discovery provides the first evidence that MDCs can form in a multicellular organism *in vivo*, highlighting the evolutionary conservation of this mitochondrial membrane remodeling process. Moreover, we show that MDC formation in *C. elegans* is triggered by refeeding following fasting, a physiologically and ecologically relevant nutritional shift. This discovery expands our understanding of mitochondrial plasticity during metabolic transitions and establishes a powerful *in vivo* model to dissect how nutrient cues shape mitochondrial membrane dynamics. By linking dietary interventions to organelle remodeling, our findings provide new insights into the role of mitochondria in maintaining organismal health and potentially modulating aging processes.

## Materials and Methods

### C. elegans strains

Worms were cultured and maintained on standard nematode growth medium (NGM) plates seeded with *E. coli* OP50-1 at 20 °C. WBM1705 was generated by crossing WBM1231 (SKI LODGE TOMM-20^aa1-49^::GFP) and WBM1688 (TIMM-50::wrmScarlet), both described previously (Valera-Alberni et al., 2024). All new strains introduced in this study are described in **Supplementary Table 1**. To make WBM1791, the *tomm-20(aa1–49)::GFP* sequence was amplified from plasmid pHW21 and knocked into the SKI LODGE cassette (*wbmIs114*) in strain WBM1356. This replaced the original *dpy-10* site with *tomm-20(aa1–49)* and added an SL2-linked *mScarlet* coding sequence under the *myo-3* promoter. WBM1301 [wbmIs107] expresses an C-terminal mScarlet-tagged HMG-5 (*C.elegans* homolog of the mammalian TFAM) under the somatic *eft-3* promoter, generated by CRISPR knock-in into the SKI LODGE cassette (*wbmIs88*) in strain WBM1214 (Silva-García et al., 2019). WBM909 was generated by CRISPR knock-in of GFP at the C-terminus of the endogenous *vdac-1* locus, using a homology repair template amplified from plasmid pAD1. To generate WBM1784, GFP was inserted at the C-terminus of wild-type *tomm-20*, resulting in the allele *tomm-20(wbm112[tomm-20::GFP])*; homozygous animals are sterile, so the strain is maintained as heterozygotes. All oligonucleotides used to generate the aforementioned strains are listed in **Supplementary Table 2**. CRISPR mixes containing the HR templates were prepared according to (Paix et al., 2015). The resulting CRISPR-edited alleles were sequence verified and then outcrossed six times to N2 before use for experiments.

### Microbe strains

OP50-1 bacteria were grown overnight in Luria-Bertani (LB) broth at 37 °C, then 100 μL of the culture was seeded onto plates and incubated at room temperature for 2 days. RNAi experiments were performed on E. coli (HT115) expressing dsRNA against the gene noted or an empty vector control. HT115 was cultured overnight in LB containing carbenicillin (100μg ml−1) and tetracycline (12.5μg ml−1). Then, 100 μl of liquid culture was seeded on NGM plates containing 100 μg/ml carbenicillin. Expression of dsRNA was induced by adding 100 μl of 100 mM isopropyl β-d-1-thiogalactopyranoside (IPTG) solution onto lawns at least 2 hours before placing the worms. RNAi constructs were obtained from the Ahringer library. RNAi studies for *miro-1/2* were performed on HT115 bacteria, whereas worms were exposed to *drp-1* or *fzo-1* RNAi grown on OP50 bacteria. Worms normally maintained on OP50-1 were transferred to HT115 bacteria by bleaching gravid adults. On day 1 of adulthood, worms were synchronized by egg lay onto plates seeded with either empty vector or the respective RNAi.

### Intermittent Fasting (IF)

Synchronized day 1 adult worms were transferred to fresh NGM plates seeded with OP50-1 (or HT115 if using RNAis) two days prior and treated with FUDR one day before the experiment. On day 2 of adulthood, worms in the intermittent fasting (IF) group were transferred to either unseeded NGM plates (fasted condition) or NGM plates seeded with OP50-1 (*ad libitum* fed condition); all plates were pre-treated with FUDR one day prior. On day 3, worms were transferred to fresh unseeded or OP50-1–seeded plates, respectively, to prevent bacterial growth on the unseeded day 2 plates. On day 4 of adulthood, worms in both the IF and control groups were transferred to NGM plates seeded with OP50-1. Worms underwent 2 cycles of IF, where 1 cycle = 2 days in the absence of bacteria followed by 2 days in the presence of bacteria. Consequently, worms in the IF group were fasted on day 4, refed on day 6, fasted again on day 8, and refed on day 10, the final day of imaging.

### Microscopy

Imaging was performed on a Yokogawa CSU-X1 spinning disk confocal system (Andor Technology) with a Nikon Ti-E inverted microscope (Nikon Instruments), using a Plan-Apochromat 100x/1.45 objective lens. Images were acquired using a Zyla cMOS camera and NIS elements software was used for acquisition parameters, shutters, filter positions and focus control. Exceptionally, the imaging of the TOMM-70::GFP ; TIMM-50:: mScarlet strain was carried out on a Zeiss Axio Observer Z1 (100x objective lens) with a LSM980 Scan Head and super-resolution images were acquired via the Airyscan detector. All images represent a single focal plane. To quantify the average structure size, the diameter of spherical mitochondrial-derived structures with a visible lumen or bright GFP-positive puncta was measured using the line tool in Fiji. The number of structures per muscle cell was quantified using the multi-point tool in Fiji. Prior to quantification, raw images were processed to enhance the resolution of GFP and mScarlet signals. A fixed ROI of 750±250 μm^2^ was selected for the muscle section in all images. Muscle sections outside this size range were excluded from analysis. Mitochondrial-derived structures were identified as large, round, GFP-positive structures that appeared brighter than the average mitochondrial tubule and excluded the mScarlet signal. Smaller, less bright, or irregularly shaped puncta were excluded from the quantification. For the quantification of the percentage of cells in Figure 1C and Supplementary Figure 1B, each data point represents a biological replicate, with 20–50 animals analyzed per condition. A muscle cell was included in the analysis if it contained at least one mitochondrial-derived structure that was enriched in GFP and excluded scarlet. For staining with TMRE (T669; Thermo Fisher Scientific), worms were placed on OP50-1 seeded NGM plates with 10 μM TMRE 24 h prior to imaging.

### Statistical analysis

Data were graphed and analyzed using GraphPad Prism 10. To quantify mitochondrial-derived structure numbers, a two-way ANOVA followed by Tukey’s multiple comparisons test was used to identify statistically significant differences between groups. For structure diameter quantification in Figure 1G, a one-way ANOVA combined with the nonparametric Kruskal–Wallis test was performed. Mitochondrial length in Fig 1B and Fig 3E was quantified using MitoMAPR (Zhang et al., 2019).

## Acknowledgments

Funding support was provided by NIH/NIA R01AG044346, R01AG067106 and R21AG056930. Fig 2D was created with BioRender. We thank the Caenorhabditis Genetics Center (CGC) for providing strains. We thank the members of the Mair laboratory and the Department of Molecular Metabolism for their valuable feedback and insightful discussions throughout the course of this project, and Pallas Yao for construction of some of the original strains. We also thank the Sabri Ülker Center for generously granting access to their spinning-disk confocal microscope. Finally, we appreciate the Micron team – Dr. Paula Montero Llopis, Dr. Praju Vikas Anekal and Dr. Adrienne Wells – at the Harvard Longwood Area for their insightful support and guidance on microscopy and experimental design.

## Author Contributions

M Valera-Alberni: conceptualization, data curation, formal analysis, investigation, methodology, project administration, validation, visualization, writing— original draft, review and editing.

A Lanjuin: methodology— *C. elegans* strain design, writing— review, and editing.

S Romero-Sanz: data curation, formal analysis, investigation, and writing— review, and editing.

K Burkewitz: methodology— *C. elegans* strain design. AL Hughes: conceptualization, methodology, writing— review and editing.

WB Mair: conceptualization, methodology, funding acquisition, resources, project administration, and writing— review, and editing.

## Supplemental information

**Supplementary Figure 1.**
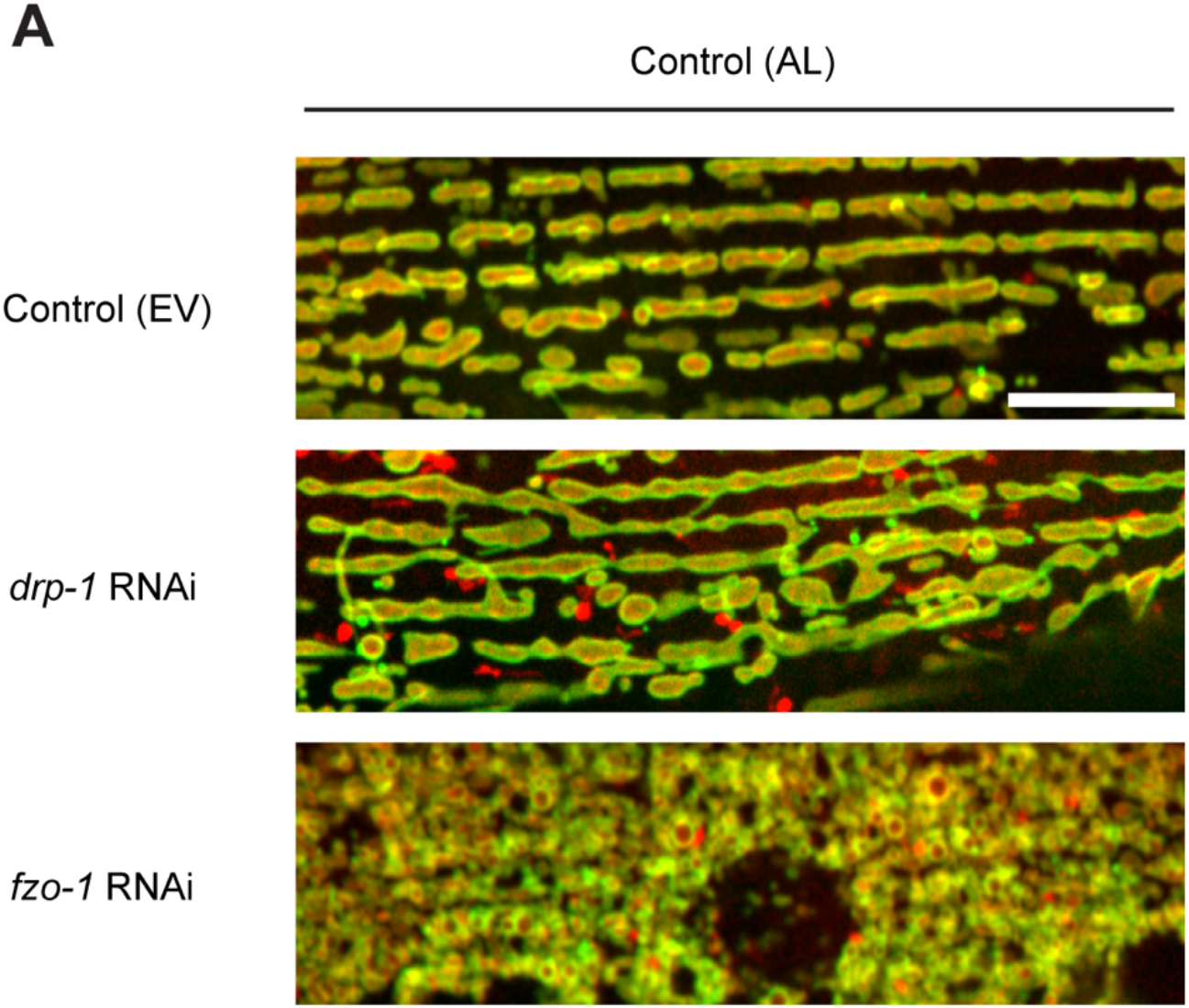
Validation of RNAi interventions. **(A)** Representative confocal fluorescence images of muscle cells from TOMM-20^aa1-49^::GFP; TIMM-50::mScarlet worms on day 6 of adulthood, maintained under *ad libitum* (AL) feeding conditions and treated with the indicated RNAi. RNAi validation shows increased mitochondrial elongation in *drp-1* RNAi-treated animals and enhanced fragmentation following *fzo-1* RNAi. Scale bar: 10 µm.

**Supplementary Figure 2.**
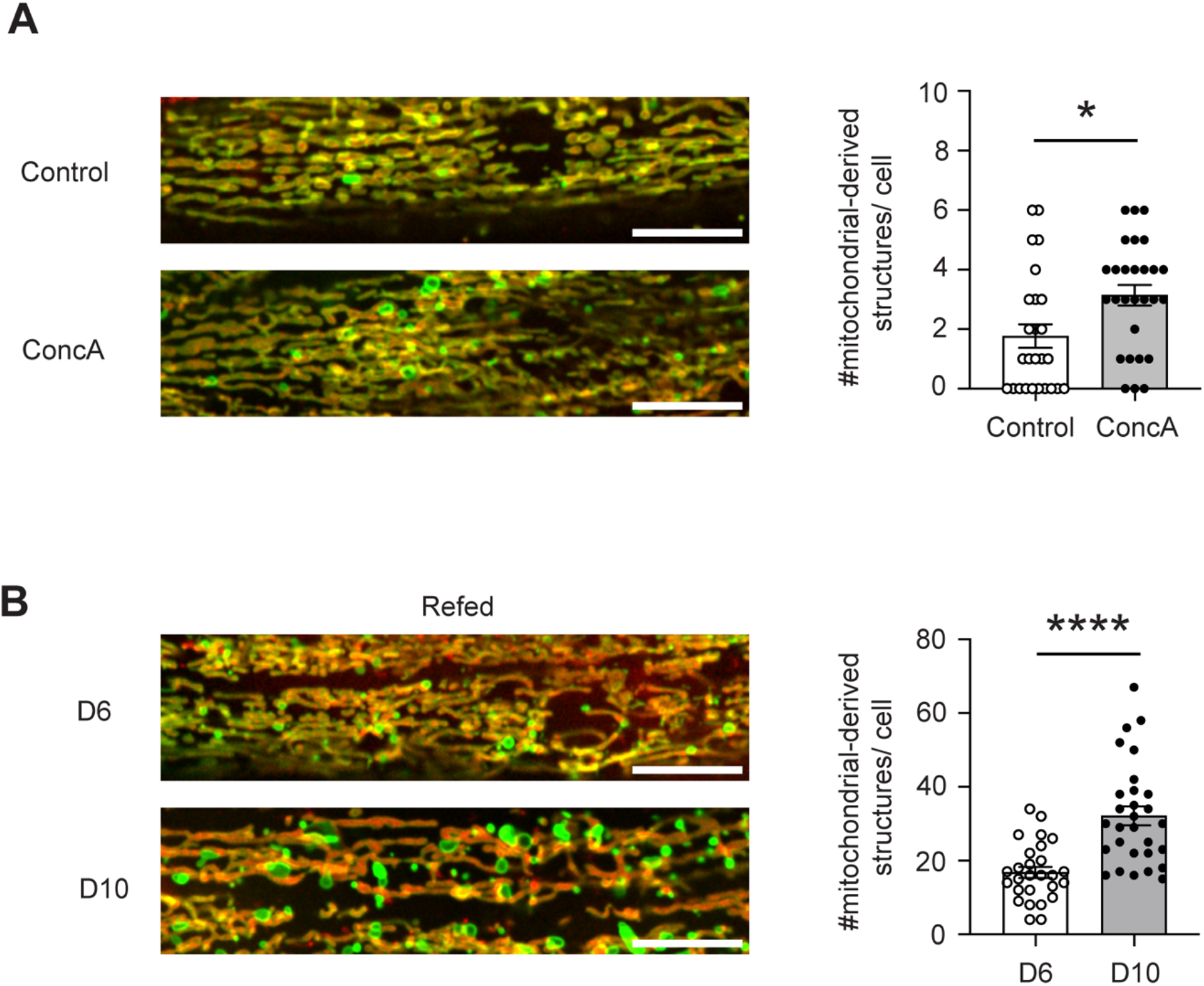
Alternative interventions promoting mitochondrial-derived structure formation in *C. elegans*. **(A)** Representative confocal fluorescence images of muscle cells from TOMM-20^aa1-49^::GFP; TIMM-50::mScarlet young adults (day 1–2) treated with Concanamycin A (2 mg/mL) for 2 hours. **(B)** Confocal fluorescence images of muscle cells from TOMM-20^aa1-49^::GFP; TIMM-50::mScarlet refed worms at day 6 (following the first fasting-refeeding cycle) and day 10 (following the second cycle). Quantification represents mean ± SEM from n = 20–30 animals per condition. *P < 0.05, ****P < 0.0001. Scale bar: 10 µm.

**Supplementary Table 1.** List of *C. elegans* strains.

| Genotype | Strain ID | Alternative name | Source |
| --- | --- | --- | --- |
| N2, wbmIs97[eft-3p::tomm-20(1-49)::GFP::unc-54 3'UTR, *wbmIs65] | WBM1231 | SKI eft-3p::tomm20(aa1-49)::GFP | CGC |
| scpl-4(wbm108[TIMM-50::scarlet]) V | WBM1688 | TIMM-50::mScarlet | CGC |
| N2, wbmIs97[eft-3p::tomm-20::GFP::unc-54 3'UTR, *wbmIs65] ; scpl-4(wbm108[TIMM-50::scarlet]) V | WBM1705 | eft-3p::tomm20(aa1-49)::GFP; TIMM-50::mScarlet | This paper |
| N2, wbmIs107[eft-3p::hmg-5::wrmScarlet::unc-54 3'UTR, *wbmIs88] | WBM1301 | TFAM(HMG-5)::mScarlet | This paper |
| tomm-20(wbm112[tomm-20::GFP])/+ V (unbalanced HETS) | WBM1784 | endogenous knock-in TOMM-20::GFP | This paper |
| N2, wbmIs164 [myo-3p::tomm-20(1-49)::GFP::SL2::scarlet::unc-54 3'UTR *wbmIs114] SKI I | WBM1791 | SKI myo-3p::tomm20(aa1-49)::GFP with cytosolic scarlet | This paper |
| vdac-1(wbm7[vdac-1::GFP]) IV | WBM909 | VDAC::GFP | This paper |

**Supplementary Table 2.**
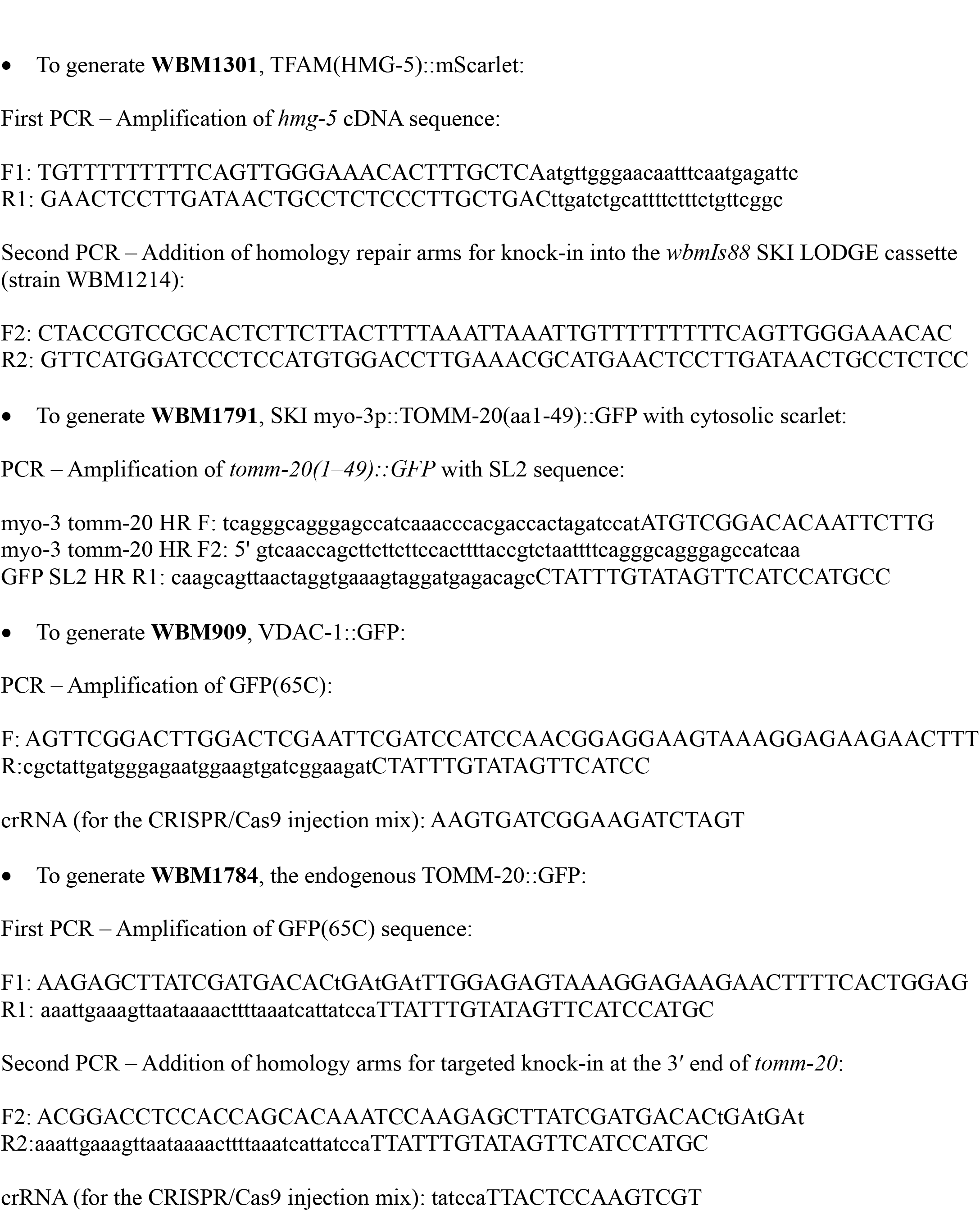
Oligonucleotides.

